# Environmental gene regulatory influence networks in rice (*Oryza sativa*):response to water deficit, high temperature and agricultural environments

**DOI:** 10.1101/042317

**Authors:** Olivia Wilkins, Christoph Hafemeister, Anne Plessis, Meisha-Marika Holloway-Phillips, Gina M. Pham, Adrienne B. Nicotra, Glenn B. Gregorio, S.V. Krishna Jagadish, Endang M. Septiningsih, Richard Bonneau, Michael Purugganan

**Author notes:** Contact Information: Richard Bonneau Michael Purugganan. these authors contributed equally to this work (Olivia Wilkins, Christoph Hafemeister). these authors are the senior and corresponding authors (Richard Bonneau, Michael Purugganan).

## Abstract

Environmental Gene Regulatory Influence Networks (EGRINs) coordinate the timing and rate of gene expression in response to environmental and developmental signals. EGRINs encompass many layers of regulation, which culminate in changes in the level of accumulated transcripts. Here we infer EGRINs for the response of five tropical Asian rice cultivars to high temperatures, water deficit, and agricultural field conditions, by systematically integrating time series transcriptome data (720 RNA-seq libraries), patterns of nucleosome-free chromatin (18 ATAC-seq libraries), and the occurrence of known cis-regulatory elements. First, we identify 5,447 putative target genes for 445 transcription factors (TFs) by connecting TFs with genes with known cis-regulatory motifs in nucleosome-free chromatin regions proximal to transcriptional start sites (TSS) of genes. We then use network component analysis to estimate the regulatory activity for these TFs from the expression of these putative target genes. Finally, we inferred an EGRIN using the estimated TFA as the regulator. The EGRIN included regulatory interactions between 4,052 target genes regulated by 113 TFs. We resolved distinct regulatory roles for members of a large TF family, including a putative regulatory connection between abiotic stress and the circadian clock, as well as specific regulatory functions for TFs in the drought response. TFA estimation using network component analysis is an effective way of incorporating multiple genome-scale measurements into network inference and that supplementing data from controlled experimental conditions with data from outdoor field conditions increases the resolution for EGRIN inference.

## INTRODUCTION

Plants alter the expression levels of different sets of genes to coordinate physiological and developmental responses to environmental changes (Nagano et al., 2012; Plessis et al., 2015). The ability to respond to environmental signals is the hallmark of adaptation and underlies tolerance to biotic and abiotic stresses. For domesticated crop species, like Asian rice (*Oryza sativa*) adaptive gene expression patterns associated with environmental changes can ensure high yields under a range of climatic conditions (Mickelbart et al., 2015; Olsen and Wendel, 2013).

Rice is a staple food for more than half of the world’s population (Khush, 2005). Changes in rice yield caused by climate change have major implications for global food security (Pachauri et al., 2014). An estimated 45% of rice growing lands are at risk of drought because they are not irrigated and depend entirely on rainfall for water (Tuong and Bouman, 2003). Moreover, many rice-growing regions have temperatures bordering critical limits for optimal grain production (Peng et al., 2004; Prasad et al., 2006; Wassmann et al., 2009). Current climate models predict marked reductions in rice yield due to changes in the frequency and intensity of extreme climate events (Pachauri et al., 2014; Redfern et al., 2012). Understanding the mechanisms that permit growth in fluctuating environments is a critical step towards identifying the molecular processes that could be targeted through traditional breeding or genetic engineering for developing stress tolerant plants.

Plants rely on gene regulatory networks to orchestrate dynamic adaptive changes that enable them to survive in growth limiting environments. Gene regulatory networks are the core information processing mechanism of the cell; they coordinate the timing and rate of genome-wide gene expression in response to environmental and developmental signals (Bonneau, 2008; Huynh-Thu and Sanguinetti, 2015; Imam et al., 2015; Roy et al., 2013; Schulz et al., 2012). Environmental gene regulatory influence networks (EGRINs) are defined by the environmentally-modulated interactions of protein transcription factors (TFs) with conserved regulatory elements in genomic DNA (Sullivan et al., 2014; Zhang et al., 2015) to effect the organization of the chromosome and the transcription of RNAs, including miRNAs and lncRNAs, that in turn have regulatory potential (Mercer and Mattick, 2013). Plant genomes encode an expanded repertoire of TFs relative to other organisms (Shiu et al., 2005) that is consistent with their sessile lifestyles and their dependence on gene response mechanisms to cope with variable environments.

The need to map out and dissect gene regulatory networks has led to the development of various experimental and computational approaches to infer their structure and composition (Bonneau, 2008; Greenfield et al., 2013; Koryachko et al., 2015; Roy et al., 2013). The simplest large scale EGRINs are based on transcriptome data, measured by high-throughput sequencing or array-based technology, without regard for other post-transcriptional and translational regulatory events that are known to influence transcriptional regulation (Koryachko et al., 2015). These methods assume that the expression of genes across environmental conditions, perturbations and genotypes can be used to predict regulatory relationships; however, many TF proteins exist in an inactive form in the cytosol or nucleus until they are activated by environmental or developmental signals (Fu et al., 2011; Ohama et al., 2016). It is not feasible to measure all of the complex and varied factors contributing to the regulation of gene expression; as such, network inference algorithms have been developed to predict regulatory interactions in the absence of complete data by incorporating additional complementary data types or prior knowledge of the network structure to estimate the effects of unmeasured regulatory layers (Arrieta-Ortiz et al., 2015; Bonneau, 2008; Fu et al., 2011; Greenfield et al., 2013; Misra and Sriram, 2013; Roy et al., 2013).

In model prokaryotic and eukaryotic systems, where much of the true architecture of many regulatory networks is known, methods that combined expression data and additional data types that define structure priors were able to infer more accurate regulatory networks that methods based on gene expression data alone (Arrieta-Ortiz et al., 2015; Fu et al., 2011; Greenfield et al., 2013). For rice, where only a minority of regulatory relationships are known, several EGRINs have been published that use additional knowledge of network regulators to inform transcriptome-based network inference models. For example, Obertello et al., (2015) included knowledge of regulatory interactions in other species, and Nigam et al., (Nigam et al., 2015) consider microRNA-mediated TF activity. In both of these instances, the additional data types were used to filter and refine co-expression networks. Sullivan *et al.* (Sullivan et al., 2014) used changes in chromatin accessibility upstream of the coding regions of genes, a hallmark of gene regulatory activity (Buenrostro et al., 2013; Zhang et al., 2015), and knowledge of cis-regulatory motifs, rather than changes in gene expression, to construct a cis-regulatory network of the response of *Arabidopsis* to high temperature and during photomorphogenesis. Mapping TFs to target genes in this manner considers putative functional elements, but does not capture the output of the regulatory network, specifically, the regulated changes in transcript abundance that expression based networks measure.

Our method combines the strengths of transcriptome and chromatin accessibility based methods in a novel algorithm that systematically incorporates multiple genome scale measurements into a single model of gene regulation. We use a combination of static (i.e. cis-regulatory motif occurrence) and dynamic (i.e. transcriptome and chromatin accessibility data) genome-scale measurements to define the proximal promoter region for all genes and to estimate the regulatory activity of transcription factors. These analyses were used as a starting point for inferring an EGRIN using an adapted version of the Inferelator, a method for learning networks based on sparse linear models (Arrieta-Ortiz et al., 2015; Bonneau et al., 2006; Greenfield et al., 2013). We incorporated time series gene expression data from controlled experiments where single environmental factors were perturbed and from agricultural field experiments collected over multiple growing seasons with experimentally induced stress responses. Our approach overcomes some of the shortcomings implicit in transcriptome based network prediction (e.g. co-expression as a proxy for regulation), by leveraging a multi-factor experimental design. Using this approach, we have assembled the first high-resolution view of global environmental gene regulation in rice.

## RESULTS

For this study, we used multiple genome scale measurements – transcriptome, nucleosome-free chromatin, and *cis*-regulatory motif occurrence – to learn the gene regulatory networks associated with response to environmental change. Briefly, our method for inferring EGRINs was as follows:(1) for each TF with known *cis*-regulatory elements (CRE), we identified target genes that had the relevant CRE in a region of open chromatin in their promoters; (2) we then used the expression of the putative target genes to estimate the Transcription Factor Activity (TFA) for each transcription factor using a method called Network Component Analysis (Liao et al., 2003); (3) we then inferred the regulatory network using the Inferelator algorithm (Arrieta-Ortiz et al., 2015; Bonneau et al., 2006; Greenfield et al., 2013), in which regulatory interactions are predicted based on (partial) correlations between TFAs and the expression of target genes.

We carried out two major experiments (Figure 1A) in which we exposed rice plants to a wide range of environmental conditions and monitored their functional and transcriptome responses over time. In the first experiment, we grew rice plants in climate controlled growth chambers for 14 days in hydroponic culture before initiating heat shock (a transfer from 30°C to 40°C) or water deficit treatments (removal of all root available water). We collected samples from the heat shock treatment every fifteen minutes for 4h; however, we collected samples from the water deficit treatment every fifteen minutes for only 2h. Beyond 2h of the water deficit treatment we began to see irreversible tissue damage, including leaf tip death, and we stopped the treatment as that response was not of interest for this study. In the second experiment, we grew three rice cultivars in two agricultural fields, one irrigated and one rain fed, during both the rainy and dry seasons and we monitored weather variables throughout the experimental period. Leaf samples were harvested every other day for 29 days during the vegetative growth phase at the same time of day with respect to sunrise. This experiment is described in detail in Plessis et al. (2015). The controlled experimental treatments differed from agricultural stresses in several critical ways:the heat shock was a near-instantaneous 10°C temperature increase and the water deficit treatment exposed the roots of hydroponically grown plants into the air. While these conditions do not replicate field conditions, they allowed us to apply consistent treatments to a large number of plants simultaneously and because they had previously been demonstrated to elicit relevant changes to both plant functional responses, and importantly for our study, to the expression levels of a number of transcripts that were *a priori* of interest.

**FIGURE 1:**
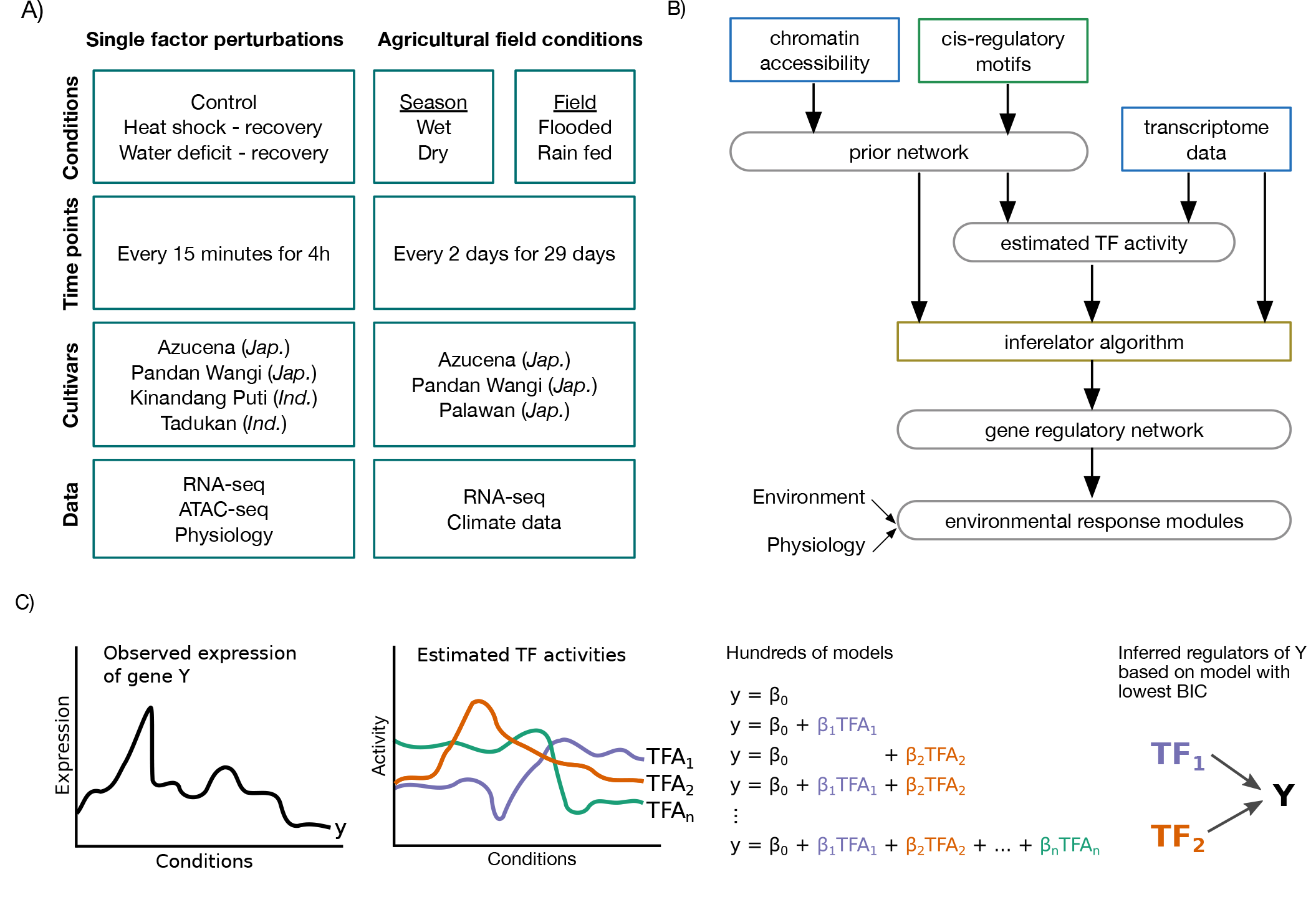
Schematic overview. A) Experimental design:We queried the response of five tropical Asian rice cultivars to high temperatures, water deficit, and agricultural field conditions. Data collected includes time series transcriptome data (RNA-seq) and chromatin accessibility (ATAC-seq) B) Data analysis:We connect transcription factors with genes via known cz’s-regulatory motifs in accessible promoters; this prior network together with the transcriptome data is used to estimate transcription factor activities; the Inferelator algorithm is used to construct the final network. C) Inferelator algorithm:For every target gene hundreds of models linking its expression to TF activities are evaluated and one model is selected to define its regulators.

We included five tropical Asian rice cultivars in these experiments, all of which were traditional land races including representatives of two of the major rice subspecies – *indica* (cultivars Kinandang Puti, and Tadukan), and *japonica* (cultivars Azucena, Pandan Wangi, and Palawan, Figure 1A). These varieties were traditionally used in either irrigated culture (Pandan Wangi and Tadukan) or in rain fed fields (Azucena, Kinandang Puti, Palawan) and their divergent ecological adaptations allow us to capture a greater breadth of responses to the environmental treatments. Our method systematically incorporates multiple genome-scale measurements, including chromatin accessibility, *cis-* regulatory motifs, and transcriptome data, to generate predictions of gene regulatory interactions in rice leaves. We interpreted the regulatory network in the context of the plant functional measurements and weather data (Figure 1B).

### Photosynthetic rate is decreased in response to environmental stress

To provide the functional context in which gene expression was measured, we monitored plant physiological status during the heat shock and water deficit treatments, including carbon assimilation rate and stomatal conductance. We observed distinct functional responses for each stress type, with similar responses observed for all four cultivars (Supplemental Figure S1) and so, hereafter, we describe the responses of Azucena as an exemplar. In response to the heat treatment, we observed an initial steep and transient decrease in the carbon assimilation rate (∼80% control) of Azucena followed by a recovery to a sustained rate of ∼90% control for the length of the heat stress treatment (Figure 2A, Supplemental Figure S1). Upon return to the lower temperature, the carbon assimilation rates remained stable, but below the rate in control conditions for the duration of the measured recovery period. The degree of change is variable across the cultivars, but that the trend is consistent across all.

**FIGURE 2:**
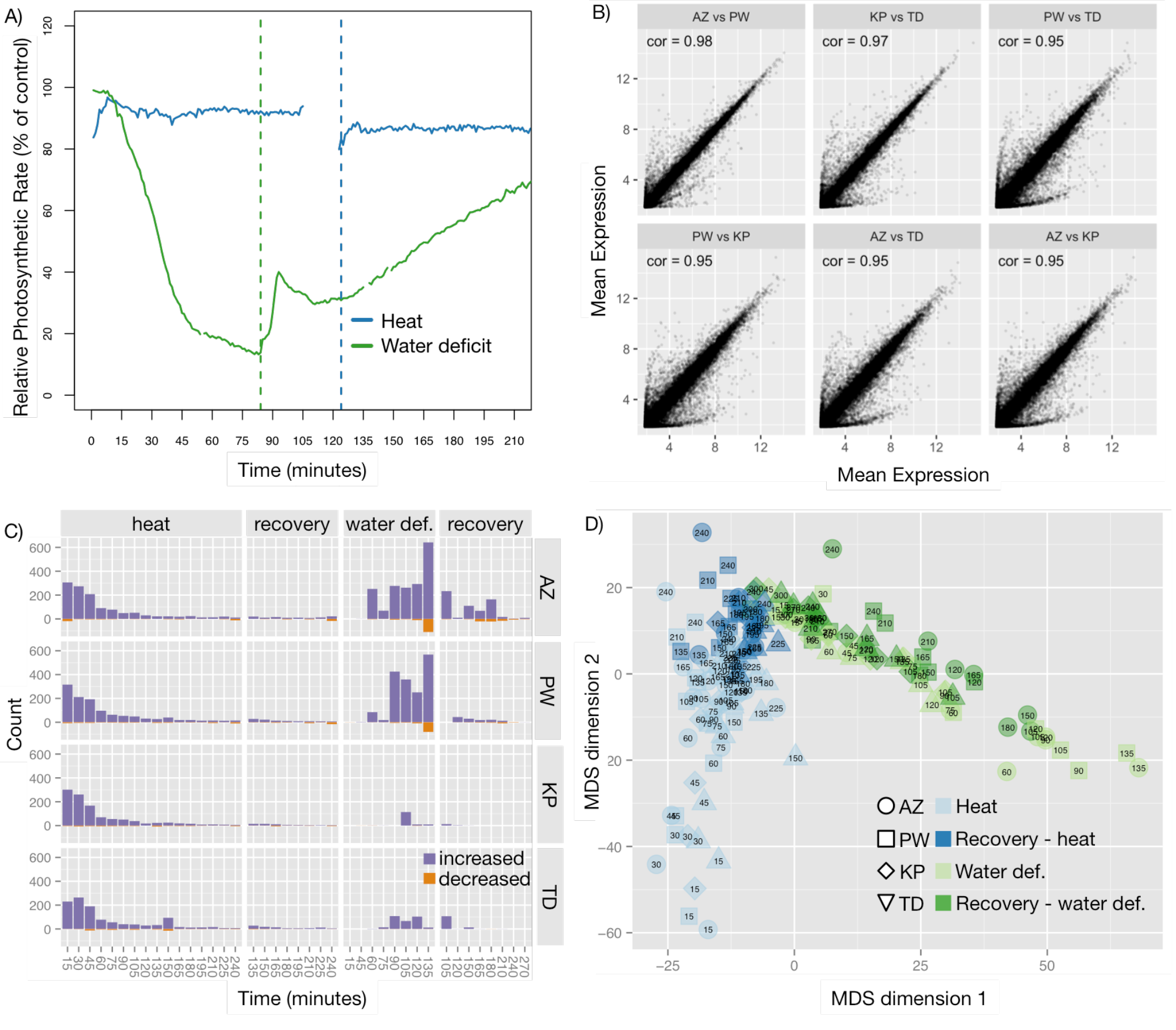
Overview of experimental data. A) Functional responses:The relative photosynthetic rate of stress treated Azucena plants is presented (n=3 for each treatment; mean of the replicates is presented). Data for other genotypes are in supplementary material (Supplementary Figure S1). The vertical dashed lines indicate the end of the stress treatment and the start of the recovery treatment. Time 0 coincides with minimum functional status observed within 15 min of the start of the stress imposition B) For each cultivar we averaged the expression across control conditions for every gene. Pairwise scatter plots and Pearson correlations for Azucena (AZ) and Pandan Wangi (PW), Kinandang Puti (KP), and Tadukan (TD) are shown. C) Number of differentially expressed genes for each genotype, time point and treatment. Genes with positive fold change are shown as positive counts in purple; genes with negative fold change are shown as negative counts in orange. D) Expression similarity of data points shown as multi-dimensional scaling plot based on Euclidean distance of log-fold-change of 2097 differentially expressed genes (see method section). The number inside each data point indicates the time in minutes since the onset of the treatment at which the sample was measured.

In response to the water deficit treatment, we observed a continuous decrease in rate of carbon assimilation over the period of the treatment. For example, Azucena’s carbon assimilation rate declines to around 90% of its well-watered rate after only 15 minutes of water deficit stress and to around 15% after 90 minutes, the last measurement before the plants were returned to the water-unlimited treatment. The other three cultivars similarly have decreasing rates of carbon assimilation over the period of the water-deficit treatment. In all four rice cultivars, we observe a gradual increase in the carbon assimilation rate when the plants are given ample water for recovery, though none of them returned to pre-treatment rates in the monitored recovery period.

For neither treatment did we observe functional differences that could be associated with the rice subspecies (*indica v. japonica*) or the type of field (irrigated *v.* rain fed) in which the rice was traditionally grown. Because the functional measures of the sampled leaves showed partial recovery upon return to the initial experimental conditions, and after four days post-treatment we did not observe leaf mortality on the stress treated plants, we are confident that the treatments did not cause irreversible damage to the sampled leaves.

### Fast responses to heat, slow responses to water deficit

A comparison of the four cultivars indicated that they had similar transcriptomes as shown by the high correlation of the gene expression between cultivars across the genome (Figure 2B, Supplemental Figure S2). The correlation of gene expression was highest within each subspecies (cor=0.98 for *japonica* cultivars; cor=0.97 for the *indica* cultivars); correlation between the transcriptomes of different subspecies was also high (cor > 0.95). Given this similarity, there should only be very low bias introduced by having aligned all cultivars to the same Nipponbare reference genome. We identified differentially expressed genes in the controlled chamber experiments in response to single-factor environmental perturbations. Results indicate that the two types of treatments (heat and water deficit) lead to responses with distinct temporal patterns of expression regulation (Figure 2C) that paralleled the leaf functional responses to stress. During the heat treatment, there is a rapid increase in the expression of ∼300 genes, but after 90 minutes only few genes remain perturbed (Figure 2C, Supplementary File S1). In contrast, the response to water deficit is much slower, with almost no differentially expressed genes detected before 60 minutes of treatment after which a large number of differentially expressed genes were observed for the remainder of the stress period. Even 90 minutes after the plants were returned to water-unlimited conditions many genes remain perturbed. The dynamics of the stress responses are summarized in a multi dimensional scaling plot based on Euclidean distance of log-fold-change of 2,097 genes that are differentially expressed in at least one condition (Figure 2D, Supplemental Figure S3). We note that, in contrast to the response to heat shock where the response of all four genotypes was similar, the response to the water deficit stress is stronger (as measured by the number of differentially expressed genes) in the Japonica cultivars (Azucena, Pandan Wangi) than in the Indica cultivars (Kinandang Puti, Tadukan); we did not observe these differences in the plant functional measurements.

Most of the differentially expressed genes in this analysis were more highly expressed in the treated than in the control conditions. The relatively small number of down-regulated genes identified in this analysis is consistent with other published abiotic stress studies in Arabidopsis (Kilian et al., 2007; Matsui et al., 2008; Seki et al., 2002) and Rice (Zhou et al., 2007) that applied severe and short term treatments, much like the one we used in this study. This imbalance is characteristic of samples collected soon after the treatment is initiated (within hours). In their 2008 study in Arabidopsis, Matsui *et* al.(Matsui et al., 2008) found that after 2h of water deprivation, a much larger number of genes were up regulated than down regulated in response to the treatment; whereas, after 10h, the number of up-and down-regulated genes was effectively equivalent. Similarly, the early time points (0.5h, 1h, 3h, 6h) in the osmotic stress treatment in the AtGenExpress Abiotic Stress Dataset(Kilian et al., 2007), had a much larger number of up-regulated than down-regulated genes; at the later time points (12h, 24h), the number of up- and down-regulated genes were approximately equal.

### ATAC-seq interrogation of rice leaves under multiple conditions reveals stable accessible regulatory regions

Nucleosome-free regions of the genome are strongly associated with active sites of transcription. We used Assay of Transposase Accessible Chromatin (ATAC)-seq to identify nucleosome-free regions of the genome (Figure 3A). According to our paired-end sequencing results the majority of DNA fragments are short, 55 bps to 65 bps long, and there is an exponential decline in the distribution of longer fragments (Supplemental Figure S4). A fragment is defined as the DNA region bounded by the forward and reverse read. To call chromosomal regions “open”, we count the number of ATAC cut sites (first base of an aligned forward read and first base after an aligned reverse read) in its proximity. We called a base open if more than half of the libraries contained at least one cut site in the 72-bp window centered on the base (Supplemental File S2). These requirements for calling open sites were chosen because of the high conservation of open sites identified across experimental conditions (Supplemental Figures S5 and S6) and because of the relatively low number of reads per ATAC-seq library which led to a low signal to noise ratio. If two open bases are fewer than 72 bps apart, we call all intermediate bases open. Based on this definition, the average open region length was 268 bps, median 206 bps (Figure 3B). In rice, the distance between nucleosome centers (dyads) ranges from ∼175 bps in promoter proximal regions to ∼191 bps in intergenic regions (Wu et al., 2014). In the human lymphoblastoid cell line in which the ATAC-seq method was developed, the fragment length distribution had a clear periodicity of ∼200 bps, with the largest peak corresponding to the length of a single nucleosome, and multiple smaller peaks corresponding to integer multiples of up to six nucleosomes (Buenrostro et al., 2013). We did not detect peaks with lengths corresponding to multiple nucleosomes in rice, which may reflect the more compact structure of plant promoters. In Arabidopsis, more than half of DNase I hypersensitivity sites within 1Kb of TSS were located within the first 200 bps upstream of the TSS (Zhang et al., 2012a), indicating that the displacement of a single nucleosome may be sufficient to permit regulated gene expression in both rice and Arabidopsis.

**FIGURE 3:**
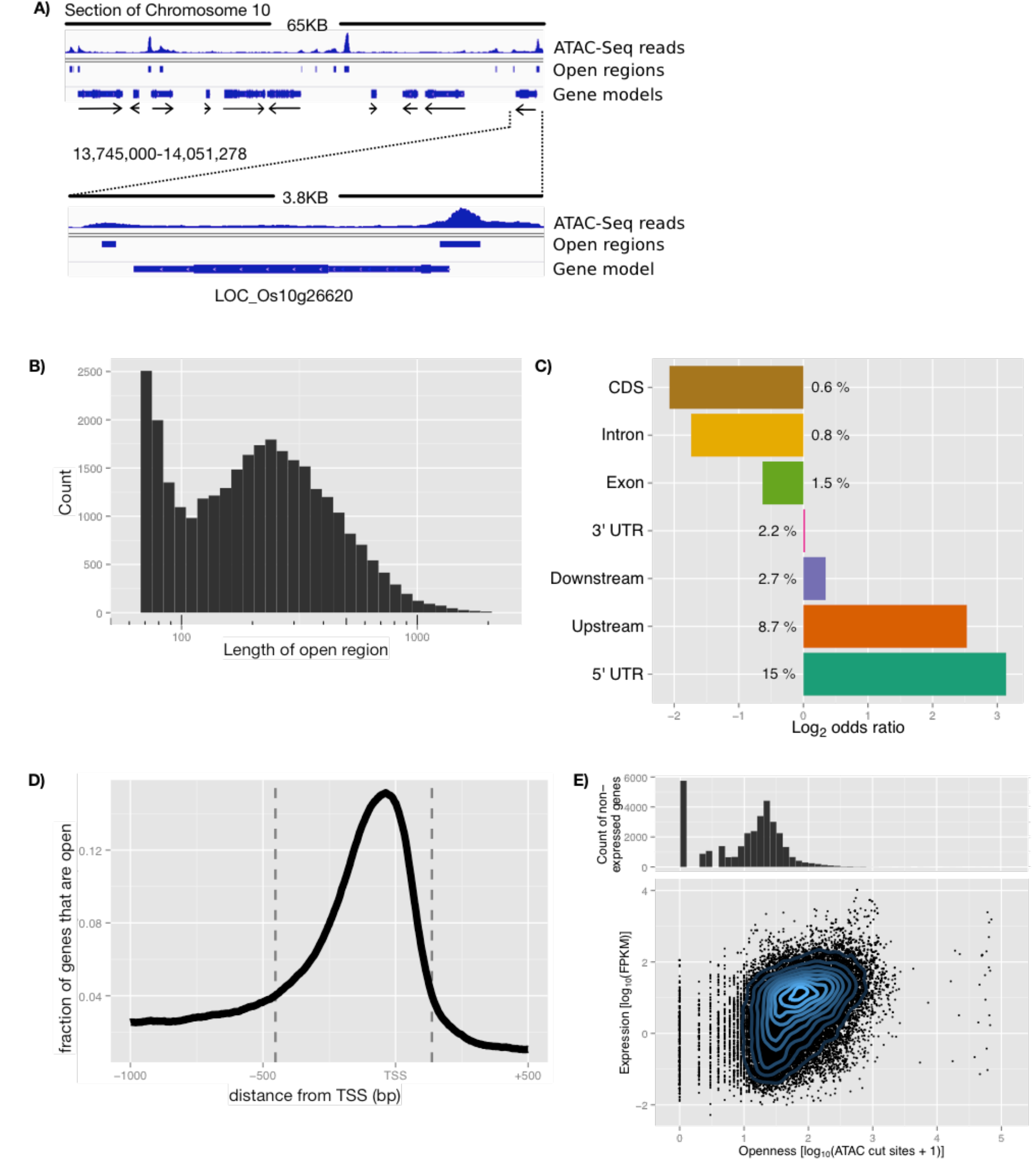
Genomic distribution of open chromatin regions identified by ATAC-seq. A) The genomic location of L0c_0s10g26620, DOF Zn-finger domain containing protein, is presented as representative ATAC-seq data. Solid black lines represent regions of genomic DNA; the histograms indicate the sequencing reads that align to a given genomic region; the blue bars indicate regions that were determined to be open; the blue bars of variable width indicate the structure of gene models; the black arrows indicate on which strand the gene is encoded. B) Length distribution of open genomic regions identified by ATAC-seq. C) Location of open chromatin regions relative to gene features, including the regions 500 bp upstream of the TSS and downstream of the 3’ end of the gene model. Numbers next to bars indicate what percentage of bases that belong to a specific feature fall into open regions. All odds ratios are highly significant (p < 1e-13). D) Distribution of open chromatin around TSSs. For all 56k genes (first isoforms only), we determined the bases that are covered by an open region in the -1000 bp to +500 bp window around the TSS. The dashed lines indicate the start (-453 bp) and end (+137 bp) of the promoter region where more than 4% of the genes are open. E) Histogram of number of ATAC-seq cuts in promoter for non-expressed genes (top). Gene expression (median FPKM across all AZ samples) vs number of ATAC-seq cuts in promoter for expressed genes; Pearson correlation 0.42 (bottom). The same y-axis is used in both panels.

Designating open regions in this manner, we identified 29,978 open regions covering ca. 8 Mbp (∼2% of the genome). The open regions were distributed through out the genome, in genic and intergenic regions, with a frequency similar to open regions identified using DNase I hypersensitivity sites (Zhang et al., 2012b) (Supplementary Figure S7). As expected nucleosome-free regions were non-randomly distributed in the genome with respect to genomic features. We examined their distribution and calculated the enrichment of bases belonging to open regions for the following features:500 bps upstream of TSSs, 5’ UTR, coding sequence, exon, intron, 3’ UTR, 500 bps downstream of gene (Figure 3C). We observed an almost 5.6 fold enrichment of bases belonging to open regions in the 500 bp upstream of the TSS of genes and an 8.5 fold enrichment in the 5’ UTRs of genes with 15% of all bases occurring in open regions (Figure 3C). Introns (∼0.3 fold) and coding sequences (∼0.2 fold) were depleted of open regions.

To get a better idea of where in the promoter region open chromatin was located for rice, we aligned all 56,000 genes in the genome with respect to their TSS. For every base from 1000 bp before the TSS to 500 bp after the TSS, we calculated the fraction of genes that were covered by an open region as described above (Figure 3D). The resulting distribution shows a sharp peak around 50 bp before the TSS where ∼15% of all genes have a region of open chromatin. The distribution quickly falls off to both sides almost reaching the background level of 2.1% at -1000 and +500 bp. We used this curve to define the promoter boundaries of -453 and +137 bp based on the coverage threshold of 4% (more than 4% of genes have open chromatin between -453 and +137 bp) as this was twice the background level. To test the effects of promoter openness on gene expression, we compared the expression of genes with the number of ATAC cut sites that fall within each gene’s promoter (Figure 3E). For this analysis we considered only Azucena growth-chamber samples and summed the ATAC results of all libraries. Of the ∼33K genes whose transcripts were not detected (FPKM of 0), 27K had 30 or fewer cut sites in their promoter. In contrast, 74% of the expressed genes had more than 30 cuts (13-fold enrichment, p < 1e-15) and we observed a significant correlation between the number of cuts and expression (Pearson correlation of 0.42).

### TF binding motif occurrence in open promoters used to derive priors on network structure

We mapped known TF binding motifs (*cis*-regulatory motifs) to open chromatin regions in the promoter regions of expressed genes to organize the existing knowledge of putative regulatory interactions. We called this collection of regulatory hypotheses the network prior. For 666 of the more than 1800 TFs in the rice genome (Jin et al., 2014), the cis-regulatory binding motifs have been determined (Matys et al., 2003; Weirauch et al., 2014). We searched for these motifs in open promoter regions defined by the ATAC-seq analysis (Figure 4A), and found significant motif matches for 445 of the TFs. By limiting the search to the open promoter regions we greatly reduced the sequence space for finding motif occurrences, which lowered the number of random motif matches. Open promoter regions made up ∼2.3 Mbp (∼0.6% of the genome) distributed among ∼9,000 genes, compared to ∼14.7 Mbp of the promoters of all expressed genes. In this way, we mapped 445 TFs to 5,447 target genes via 77,071 interactions. The median out-degree (the number of regulatory edges initiating from a TF) of TFs with targets was 75; the median in-degree of targets with TFs was 8. For example, the known binding site of Heat Shock Factor A2a (OsHSFA2a - LOC_Os03g53340) is the Heat Shock Element (HSE). Fifty-three expressed genes had the HSE in an open region of their promoter, and were mapped as targets of OsHSFA2a in the open-chromatin network prior (Figure 4B).

**FIGURE 4:**
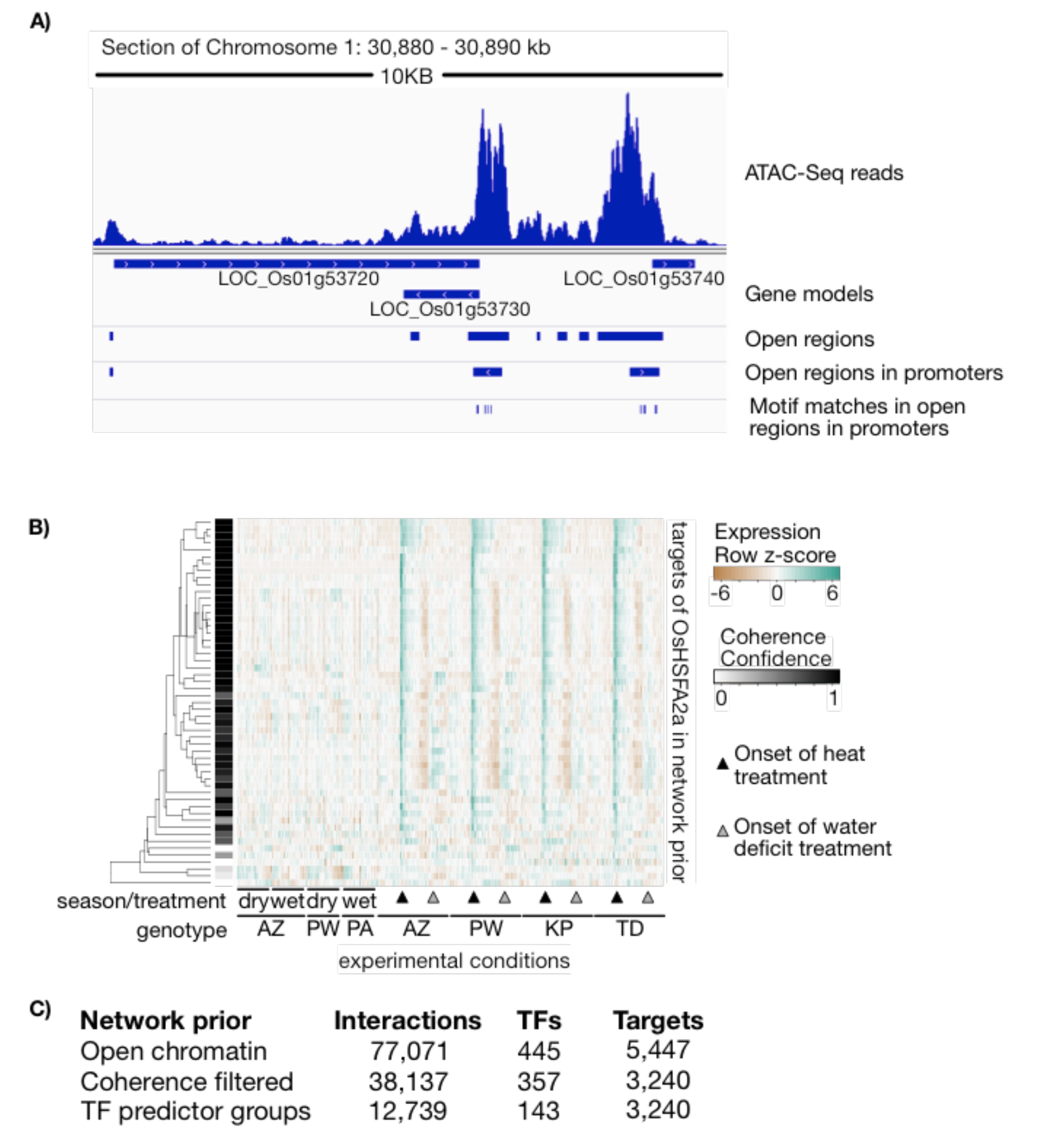
Prior knowledge of the network structure is used to calculate TF Activity. A) Exemplar of network prior inputs. Region of chromosome 1 encoding 3 genes is highlighted to indicate how motif locations gave rise to the network prior. B) Coherence filtering of network prior, showing OsHSFA2a as an example. Heatmap of expression of network prior targets with higher levels of expression indicated with green tones, and lower expression levels indicated with brown tones. Coherence scores are indicated with grey scale bar – darker tones indicate higher confidence. Sample order details available in Supplementary File S3. C) Number of regulators, edges and targets in each stage of the network prior.

We used the expression data to identify interactions in the network prior that were likely to be regulatory, based on coherent expression profiles. This step was critical, as a substantial fraction of interactions in the network prior was not expected to be related to regulatory TF binding in our experimental conditions. To identify and remove nonfunctional interactions from the network prior, we assumed that for a given TF, true prior targets should show coordinated expression across at least some of our experimental conditions, and that the targets of the TF should be enriched for that particular expression pattern with respect to background. An example output of this step for the OsHSFA2a is shown in Figure 4B, sample order for this heatmap and all other plots is given in Supplemental File S3. Each target gene was given a score indicating our confidence that it was a true target of the TF based on the criteria described above (see Method section for details); genes with low scores were removed from the network prior. This coherence-filtered network prior had 38,137 interactions (a reduction of 51%); 357 TFs were connected with 3,240 target genes (Figure 4C, Supplemental File S4). Of the TFs with targets, the median out-degree was 44; of the targets with TFs, the median in-degree was 5. Some TFs had identical target gene sets in the network prior. This occurred in cases where the TFs were members of large TF families with identical DNA binding motifs. For example, there are 25 HSFs in the rice genome, all of which are predicted to bind to the same HSE. As a consequence, all HSFs have identical targets in the prior network; we group these TFs together as a TF predictor group. A total of 276 TFs formed 62 TF predictor groups (Supplemental File S5). This generated a final network prior of 13,937 interactions, connecting 143 regulators (individual TFs and predictor groups) with 3,240 target genes (Figure 4C).

### Transcription factor activity estimation from known regulatory targets

The regulatory activity of a TF can be derived from the changes in the expression of its target genes across experimental conditions. Our method estimates TF activities (TFAs) based on partial knowledge of TF-target relationships – regulatory interactions in the coherence filtered network prior - with the aim of then using the TFAs to learn better TF-target relationships. Based on the prior network and the expression data, we used network component analysis (NCA) (Liao et al., 2003) to estimate TFAs. The principal idea of NCA is to use a simplified model of transcriptional regulation and treat the known or putative targets of a TF as reporters of its activity. This approach requires at least some known TF targets, hence we estimated the activities only for the 143 regulators with targets in our network prior (Supplementary File S6). Many TFs had similar TFAs in our experimental conditions that could be associated with the different experimental treatments (Figure 5A). The activities indicate that some TFs primarily regulate their target genes in response to only one of the experimental treatments (i.e. heat or water deficit) while others regulate the expression of their targets in response to multiple experimental treatments (e.g. heat and water deficit). Moreover, we identify TFs with similar activities in the controlled experiments, but with divergent responses in the agricultural setting.

**FIGURE 5:**
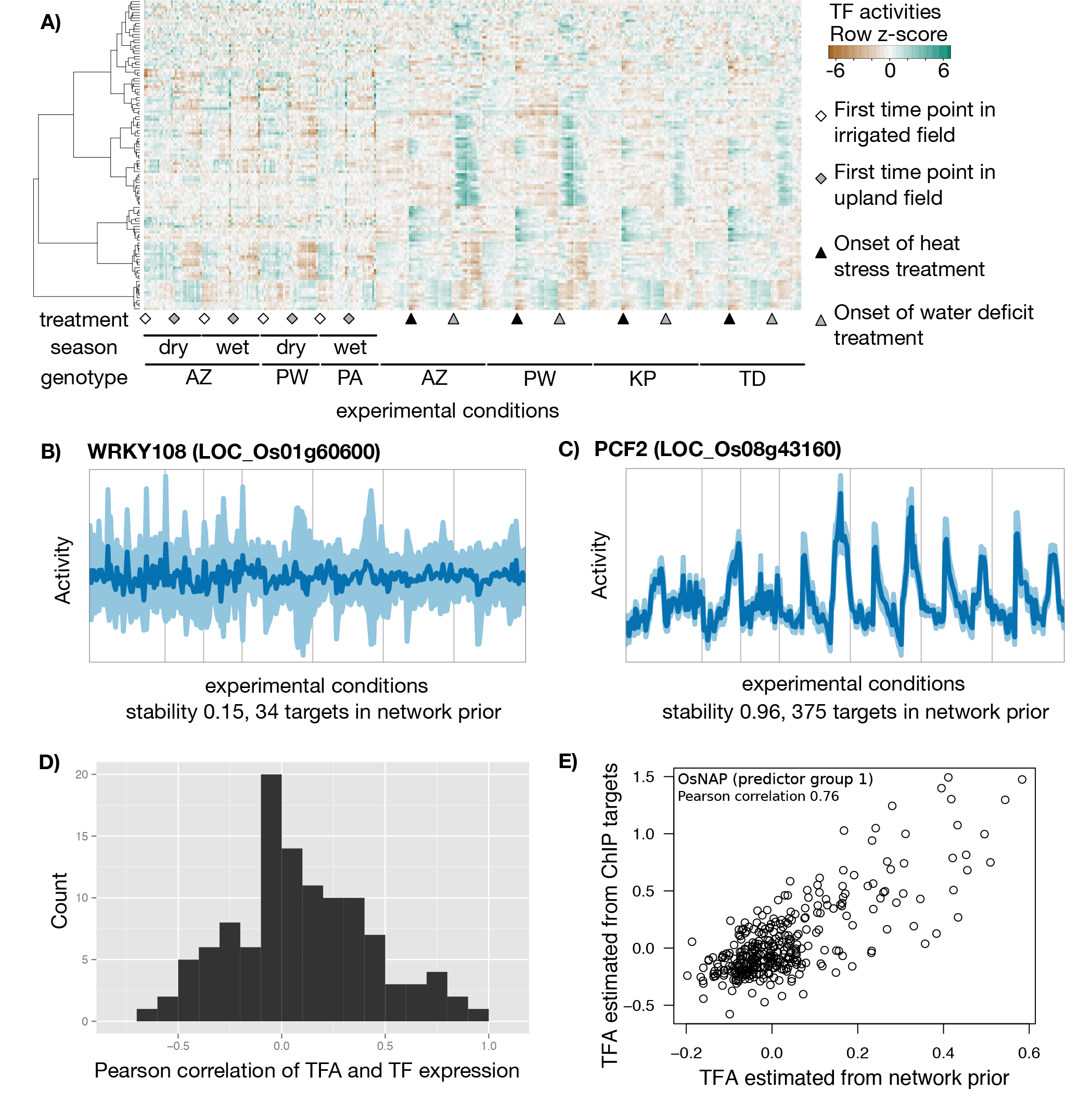
Transcription Factor Activities. Sample order is maintained for all heatmaps and TFA plots (Supplementary File S3). A) Heatmap of TFAs for all TFs and TF predictor groups for which activities were calculated. Higher levels of expression are indicated with green tones, and lower expression levels are indicated with brown tones. B) TFA stability (quantified as correlation of TFA estimates between 201 bootstrap subsamples of the network prior) for LOC_Os01g60600 and C) LOC_Os08g41360. In these plots, the light band shows the region between the 5th and 95th percentile of TFAs; the dark line shows the average TFA. D) Pearson correlation between expression and TFA for each TF. E) Correlation of TFA for OsNAP when estimated from network prior (x-axis), or from ChIP-PCR targets (y-axis).

Because the network prior was constructed from predicted TF-target interactions based on *cis*-regulatory motifs determined by *in vitro* TF binding to protein binding microarrays we anticipated that it could contain many incorrect or irrelevant interactions. Therefore, we investigated the impact that simulated changes in the network prior had on the estimated TFAs. To do so, we subsampled the prior matrix with replacement (keeping only 63% of the interactions on average) 201 times. In this way we obtained 201 TFA estimates for every TF; we called the mean pairwise Pearson correlation ‘TFA stability’ (Supplemental Figure S8, Supplemental File S7). TFA stability ranged from 0.15 (Figure 5B) to greater than 0.96 (Figure 5C). As expected, predictors (TFs and TF predictor groups) with few targets in the network prior (fewer than eight) show low stability. In the remaining set of predictors, we see 64 (52%) with very stable activities (> 0.75), which shows that at least parts of the prior network are self-consistent and estimated activities for those TFs are systematic rather random. For TFs with greater than four targets, there does not appear to be a relationship between the number of targets and stability score (Supplementary Figure S8). We found that for the majority of TFs activities and expression profiles were poorly correlated, with 85% of correlation values between -0.5 and 0.5 (Figure 5D), consistent with the fact that posttranscriptional and posttranslational mechanisms are important for TF activity and localization.

Very few validated regulatory interactions exist for rice; thereby, making it impossible to benchmark accuracy of the estimated TFAs. For OsNAP (LOC_Os03g21060), a transcription factor associated with chlorophyll degradation, nutrient transport and other senescence-related genes, seven direct regulatory targets have been experimentally validated by chromatin immunoprecipitation (ChIP) PCR (Liang et al., 2014). We estimated the TFA of OsNAP from the ChIP-validated regulatory targets (regulatory targets from (Liang et al., 2014) and expression data from the present experiment) and observed a 0.77 correlation with the TFA estimated from our distinct network prior (Figure 5E). While this result increases our confidence in our estimated TFA for OsNAP, there is not sufficient ChIP data available to generalize the concordance between our method for predicting regulatory targets and the gold-standard ChIP data.

### EGRIN:a dynamic model of transcriptional regulation in response to environmental changes

In a second step, we used the observed gene expression and the estimated TF activities to infer the EGRIN (Figure 1C). Inference is based on the Inferelator algorithm which was most recently used to infer an experimentally supported model of the *Bacillus subtilis* transcriptional network (Arrieta-Ortiz et al., 2015). The underlying idea of the method is to model the observed expression of every gene as a linear combination of the activities of a small number of TFs regulating it. For a given target gene, we constructed all models that corresponded to the inclusion and exclusion of TFs from the network prior in addition to those TFs that show a high mutual information with the target. From this large set of models, we select the one with the lowest Bayesian Information Criterion (implementing a trade-off between model complexity and goodness of fit) while slightly favoring models that also agree with the network prior (Greenfield et al., 2013) (see Method section). The inferred network is not a coexpression network, but a network where regulatory interactions are inferred from the activity of transcription factors (estimated from the expression of a subset of putative target genes) and the transcript profiles of target genes. We have evaluated this method in *Bacillus subtilis* and yeast and have shown that it is robust to false interactions in the prior network and that the inferred network was consistent with genome-wide expression levels following TF knock-out (Arrieta-Ortiz et al., 2015).

To estimate the error incurred by our EGRIN inference and to better rank regulatory interactions, we bootstrapped the expression data by subsampling from the conditions with replacement. Additionally, to control for the observed variability in TFAs with respect to small changes in our prior network (i.e. TFA stability), we also subsampled the network prior at every bootstrap as described above. Our approach then only chose interactions for the final consensus network if the predictor had a stable activity that predicted the target expression across a broad range of conditions. Given multiple bootstraps we kept those interactions that were present in more than 50% of the bootstrap networks. TFs with low stability were less likely to recall the same regulatory interaction in a large number of the bootstraps and as a consequence were less likely to be assigned target genes in the consensus network. As we increased the number of bootstraps, the network converges to a core set of interactions quickly — after 150 bootstraps more than 98% of the interactions remain stable after adding more bootstraps (Supplemental Figure S9). To obtain our final rice EGRIN, we performed 201 bootstraps. The final EGRIN contains 4,151 nodes (TFs, TF predictor groups and target genes) and 4,498 interactions (Figure 6A, Supplemental Files S8 and S9). Of all predicted interactions, 18% (796) were also in the network prior. TFs regulated between 1 and 355 target genes (Figure 6B); targets genes were regulated by 1 to 3 TFs. TFs that were grouped in the network prior because they had identical targets were included individually in the network inference as potential target genes, but only as one potential regulator.

**FIGURE 6:**
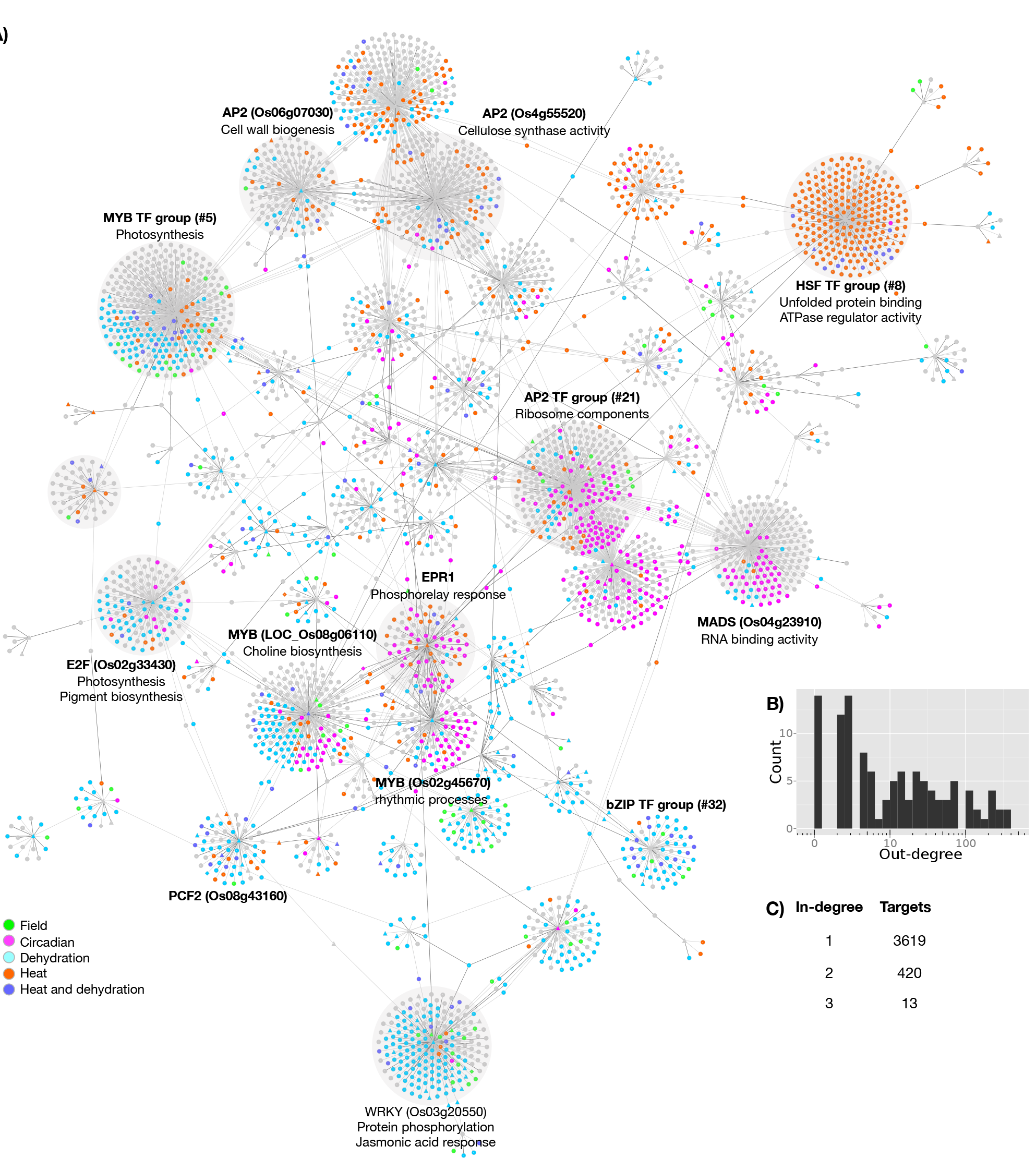
Gene regulatory network for rice leaves across environmental conditions. A) The inferred network. TFs are noted with triangles; other genes are noted with circles. The colours indicate the conditions in which the genes were differentially expressed. B) Histogram of TF out-degree frequency. C) Table of target in-degree frequency.

### Known and novel regulatory control of coordinated biological processes

The predicted rice regulatory network consists of 113 TFs, around half of those (62) are TF predictor groups that were created by grouping TFs with identical targets in the prior. In total, 4,498 interactions connect the TFs to 4,052 genes. The number of targets for each regulator, and the number of regulators per target genes are shown in Figure 5B and 5C, respectively. Fifteen predictors (11 TFs and 4 TF predictor groups) had more than 100 targets in the EGRIN. For these 15 predictors, 62% to 96% of the predicted targets were novel, i.e. not in the network prior. Many of the genes in the network were differentially expressed in response to some of our experimental treatments. We marked groups of genes on the inferred network (Figure 6A) based on the treatments in which they were differentially expressed, including heat shock and water deficit. We also marked genes as ‘circadian’ if time of day was a good predictor of their expression in the chamber experiments or as ‘field’ if the field conditions contributed more to the overall variance than the chamber data. This notation reveals that most TFs and TF groups regulate target genes primarily in response to one treatment, while a small number of TFs regulate targets in multiple conditions. For example, TF group 5, which includes two MYB TFs (LOC_Os01g09640 and LOC_Os05g10690), had the largest number of targets of any regulator in the inferred network (355 target genes). These targets are enriched for genes involved in photosynthesis, and many of them are differentially expressed in response to both the heat and water deficit treatments – stresses that altered the photosynthetic rate in our experimental conditions (Figure 2A).

We performed Gene Ontology (GO) term enrichment analyses on the target genes of each regulator and found that the targets of 27 regulators in the network were functionally enriched for a biological process or molecular function (each p < 0.0001, Supplemental File S10). Based on these analyses, we identified regions of the network whose genes were enriched for particular functions and which respond to particular environmental signals. For example, we identified a region of the network that was enriched for genes involved in RNA binding and for ribosomal structural components whose expression varied primarily as a function of the time of day. We also identified regions of the network enriched for genes with functions associated with kinases and transporter functions whose expression varied primarily in response to water deficit. Finally, we also identified regions of the network that were enriched for genes associated with photosynthesis, cell wall biosynthesis, and cellulose synthase functions whose expression vary in response to diverse environmental stimuli.

### Recapitulating and extending the known functions of HSFs

Resolving large families of transcription factors with similar binding sites is a critical problem in genome-wide regulatory studies. Here we initially grouped TFs with identical binding sites, and thus identical targets in the network prior, because their estimated activities were identical following the first step in our procedure (TFA estimation). Although we learned a regulatory model for the control of each TF in large TF groups separately, we had limited resolution to distinguish different outgoing edges for these large TF-family members (the in-degree in our network is an individual TF property and the out degree is instead an aggregate property of the members of the TF group). This limitation suggests that future experiments aimed at investigating TFs in large families with nearly identical binding sites are needed. Thus for the largest TF protein-families we lack the resolution to determine which of the TFs in the predictor group are regulating the expression of the target genes.

For the most well studied predictor group in our network, the 25 Heat Shock Factors (HSFs), we wanted to determine if *post hoc* we could identify which of the HSFs were the most likely regulators of the inferred target genes. All HSFs were connected to 46 potential target genes in the network prior via the canonical Heat Shock Element (HSE) TTCnnGAAnnTTC. The estimated TFA for the predictor group increases rapidly after the onset of the heat shock treatment, peaking after 30 minutes and then slowly decreasing over the course of the stress period; upon return to the pre-stress temperature, there is a rapid decrease in TFA – lower than the activity in control conditions – that persists for the remainder of the measured time points (Figure 7A top panel). In the EGRIN, the HSF predictor group regulates the expression of 240 target genes. The targets include a number of genes involved in the unfolded protein response, including Heat Shock Proteins (HSPs) – HSP70, HSP110, DnaJ, and DnaK - and other classes of chaperone proteins (T-complex proteins, chaperonins etc.).

**FIGURE 7:**
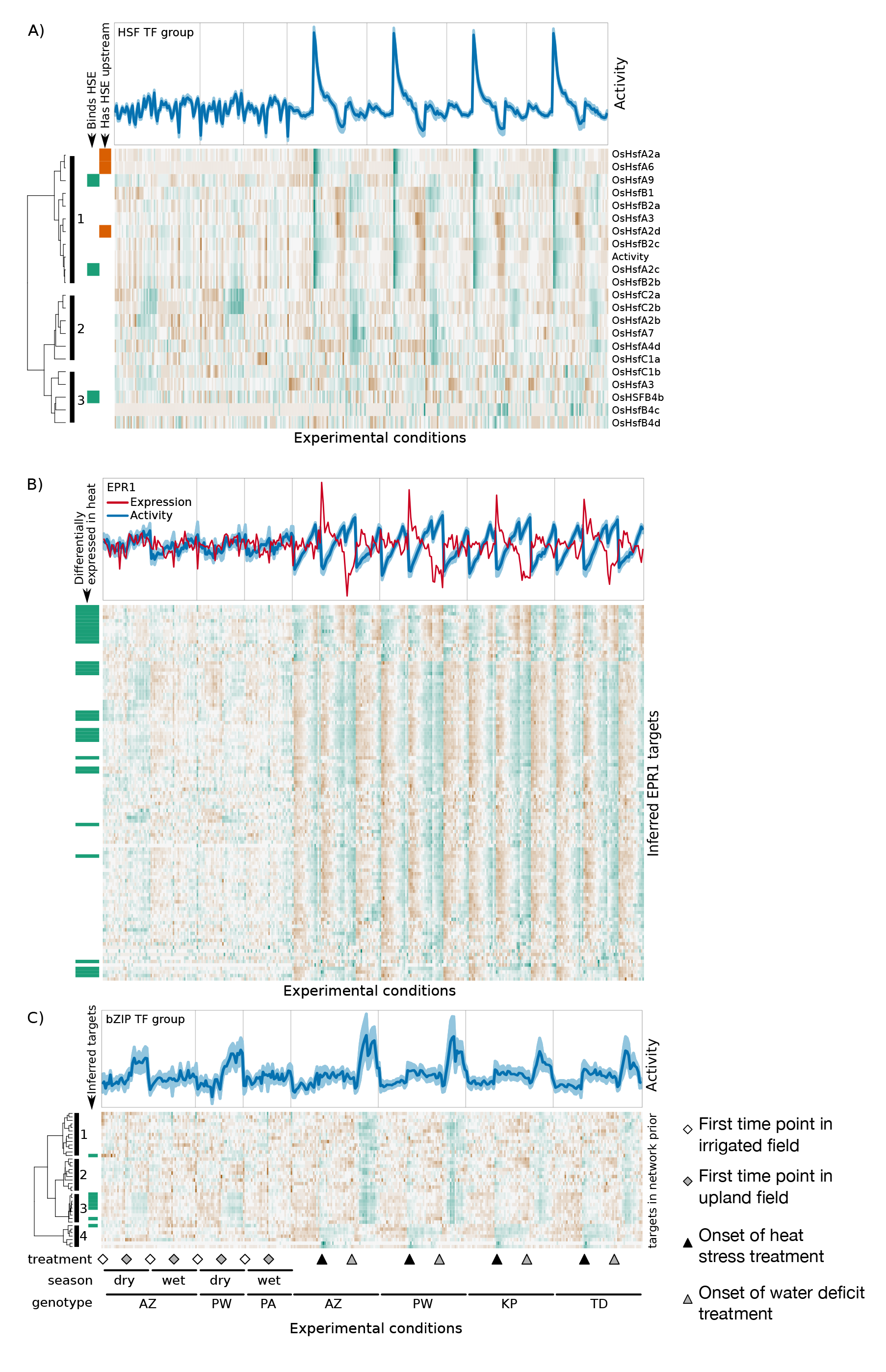
Post-hoc interpretation of inferred regulatory relationships. In all TFA plots, the light band shows the region between the 5th and 95th percentile of TFAs; the dark line shows the average TFA. In all heatmaps, higher levels of expression are indicated with green tones, and lower expression levels are indicated with brown tones. Sample order is maintained in all TFA and heatmap figures (Supplementary File S3). A) TFA of the HSF predictor group (top). Heatmap of TF transcript abundance of all group members (bottom). Subgroups are noted to the left of the figure. B) TFA (blue) and transcript abundance (red) of EPR1 (top). Heatmap of abundance of inferred EPR1 targets. C) TFA of bZIP predictor group (top). Heatmap of transcript abundance of targets in network prior (bottom).

While our method is based on the assumption that the expression of a TF will often not be indicative of its activity, TFs whose expression is highly correlated with the group’s activity are prime candidates for being the true regulators of the group’s targets. We therefore examined the expression profiles of the HSFs and compared them to the TFA for the HSF predictor group (Figure 7A bottom panel). The transcripts of four HSFs were undetectable in all of our samples; the remaining 21 formed three distinct subgroups based on their expression profiles:HSFs in subgroup one were characterized by increased expression in response to heat and water deficit; HSFs in subgroup two were primarily induced in response to water deficit; and, the expression of the HSFs in subgroup three was not clearly associated with any experimental treatment. The TFA for the HSF predictor group was embedded in subgroup one, suggesting that the true predictor could be found amongst them.

Seven of the ten HSFs in subgroup one are targets of the HSF predictor group in the EGRIN. Three of them have the canonical HSE in their promoter regions, as defined above. OsHSFA2d and OsHSFA6 each have one upstream HSE, and OsHSFA2a has three, indicating that they are potentially regulatory targets of HSFs in addition to their potential role as regulators. The other five HSFs in this group (O*s*HSFA2c, O*s*HSFA4b, O*s*HSFB2b, O*s*HSFB2c, O*s*HSFA9) do not have HSE in their promoters, and so are likely not direct targets of HSFs. We reasoned that these HSFs might be the regulators of the inferred target genes of the HSF predictor group. O*s*HSFA2c is the TF with the highest correlation between transcript abundance and TFA in our entire data set (cor=0.92). Moreover, the binding of O*s*HSFA2c is temperature regulated *in vitro* (Mittal et al., 2011) supporting its role as a heat responsive transcriptional regulator acting through the HSE.

The over-expression of O*s*HSFA7, a member of HSF subgroup two, enhances growth and survival in response to salt stress and water deficit by unknown mechanisms (Liu et al., 2013). Consistent with a role in the water deficit response, we find that the expression of all HSFs in subgroup two were induced in response to water deficit. However, the TFA, which is based on the occurrence of the HSE in open promoters, appears to be exclusively responsive to the high temperature treatment in our data set. We therefore hypothesize that the role of HSFs as regulators of the water deficit response are mediated by an unknown regulatory element other than the canonical HSE.

### Dissimilar activity and expression hint at novel functional roles of TFs

The estimated activity of a TF is not typically correlated with the expression of that TF itself; it depends only on the expression of its targets in the network prior and which other regulators are assigned to those targets. As a result, we observe a wide range of relationships between TF expression and TFA specific to each TF. One interesting example is Early Phytochrome Responsive 1 (EPR1, LOC_Os06g51260), a MYB gene that is a core component of the circadian oscillator (Filichkin et al., 2011) that binds to the canonical Evening Element, AAAATATCT, *in vitro* (Weirauch et al., 2014). In our data, we estimate that the activity for EPR1 increases over the experimental period in the growth chamber – a 4.5h period – regardless of the experimental treatments across all four genotypes, suggesting that most of its 114 target genes in the network prior were under circadian control as expected (Figure 7B top panel). We examined the expression of these target genes in an independent data set that was designed to monitor gene expression at many time points throughout the day in rice (Nagano et al., 2012) and we found that their expression was indeed oscillating daily (Supplemental Figure S10). In our inferred network, EPR1 is the regulator of 107 target genes, of which 44 were also in the network prior (Figure 6B bottom panel).

The expression of EPR1 is poorly correlated with its estimated TFA (cor=-0.52). In the control and water deficit treatments, the levels of EPR1 expression are relatively unchanging. However, at the onset of the heat stress treatment, there is a rapid and transient increase in EPR1 expression, which returns to the levels of the control condition for the remainder of the heat stress treatment; upon return to the pre-stress temperature, there is a steep and transient repression of EPR1 expression which returns to the level of the control conditions over the course of the recovery period. This expression profile is consistent with the activity of the HSF predictor group, and in fact, EPR1 is a target of the HSF predictor group both in the network prior and in the inferred regulatory network. In Arabidopsis, *At*HSFB2b binds to the promoter of another circadian clock component, Pseudoresponse Regulator 7 at a canonical HSE and is required for maintaining accurate circadian clock rhythms in high temperature and salt stress conditions (Kolmos et al.,2014). Notably, the better-characterized EPR1 ortholog in *Arabidopsis thaliana* also responds to heat stress (Kilian et al., 2007), though its role as a regulator of response to heat has not been investigated. These findings suggest that EPR1 may be an entry point for integrating the heat stress response with the circadian clock. The coordination of stress responses with the circadian clock is thought to be an adaptive strategy to maintain plant growth during periods of stress (Seo and Mas, 2015; Wilkins et al., 2009, 2010)

### Dynamic agricultural environments increase resolution of EGRIN

Upon closer examination of the network prior and the inferred network, we found evidence of the additional value of combining growth chamber and field experiments. Because these experiments perturbed different parts of the regulatory network (e.g. different time scales and treatment duration, complexity of environments, age of plants) we were able to expand the network beyond what simply increasing sample size for any one of the two experimental designs would have afforded. By combining them in a single analysis we were able to resolve parts of the network where target genes expression was similar in response to some environmental conditions, but quite different in response others. The bZIP predictor group (TF predictor group 32 in supplementary material) is one example of a predictor where targets in the network prior showed mostly correlated expression in growth chamber data but could be divided into finer groups based on the field data (Figure 7C). Prior target groups 1, 2, and 3 show distinct expression patterns only in the field; the expression of target group 3 is most similar to the estimated activity of this predictor group. The group is composed of three bZIP TFs (LOC_Os02g49560, LOC_Os08g38020, LOC_Os09g29820) that bind to a five-nucleotide motif CACGT. The estimated TFA for this predictor group shows increasing activity in response to the water deficit treatment in the growth chambers and then decreasing activity in the drought recovery period; moreover, based on the data collected in field, the predictor group’s activity increases over the course of the dry season only in the rain fed fields (Figure 7C). We infer 66 targets of this predictor group, including a number of genes involved in cellular protection in response to water deficit:trehalose-6-phospate, three Late Embryogenesis Abundant proteins involved in osmotic stress response, five dehydrin proteins, and three protein phosphatase 2C genes.

## DISCUSSION

The aim of this work was to reconstruct an environmental gene regulatory interaction network in rice leaves in response to two of the most important environmental stresses that impact agricultural productivity – high temperature and water deficit. For this purpose, we generated two genome wide data sets:1) 720 RNA-seq libraries generated from plants grown in heat and water deficit stress experiments in controlled environments and from plants grown in agricultural field conditions, and 2) the first ATAC-seq data set that identifies open chromatin sites in rice. We combined these data types with the largest available collection of TF binding motifs to generate a network of regulatory hypotheses for 4052 genes regulated by 113 TFs and TF groups.

We have previously published our method for incorporating prior knowledge into network inference using Network Component Analysis and the Inferelator algorithm for the prokaryotes *E. coli* and *Bacillus subtilis* (Greenfield and Hafemeister *et al.,* 2014; Ortiz and Hafemeister *et al.,* 2015). The present manuscript is the first use of this method to a multicellular eukaryotic organism and differs from our previous projects in several important ways:the rice EGRIN used novel data types (ATAC-seq *cis*-regulatory motif occurrence) and lower confidence interactions for learning the network prior; moreover, this was the first use of these methods on a eukaryotic organism with a complex genome. In previous publications, many of the true regulatory interactions in the network were known, and the network prior consisted of *bona fide* regulatory edges, determined using transcription factor binding assays (e.g. ChIP-seq) and genetic lesions. In the present analyses, we developed the network prior by combining two genome-scale measurements – known and predicted *cis*-regulatory motif occurrence with a map of transposase accessible chromatin – neither of which directly measure the interaction between a protein transcription factor with its cis-regulatory element in. We also developed new methods for filtering out low confidence edges from the prior network, as well as implementing a more rigorous sampling scheme (bootstrapping the expression data and the network prior) to account for the uncertainty in prior interactions.

The final inferred network includes regulatory edges that were not present in the network prior, and discounts many “false positive” edges from the network prior. So, while the network prior was based on ATAC-seq and motif occurrence data, the final inferred network was based on estimated TFA and transcriptome (RNA-seq) data. The regulatory edges present in these networks had limited overlap. It was important to permit the network inference algorithm to learn novel regulatory edges and to exclude some edges in the network prior for several reasons. We expect that the network prior would be incomplete because, we do not believe that our ATAC-seq data saturated the open sites in the genome (ie. more open chromatin may exist than we were able to identify in this experiment), we anticipate that some relevant cis-regulatory motifs occur outside of the promoter region we defined, and many TFs were not associated with known *cis-* regulatory motifs and so were excluded from our analyses. Moreover, we expect that false positive edges (connections between TFs and target genes in the network prior that are not truly regulatory) will exist in the network prior because not every *cis*-regulatory motifs occurence has a regulatory role in the genome. By increasing the number of reads in each ATAC-seq library, it may be possible to identify TF footprints in open chromatin regions (Buenrostro *et al.* 2013). In this case, the network prior could be defined with greater precision. However, it is unclear what the overall consequences this would have on the estimation of TFA or on the inference of the regulatory network.

One of the challenges of engineering agricultural crops with improved stress tolerance is translating experimental advances from controlled conditions in the growth cabinet into the complex and fluctuating environments found in the field. The disjunction between plant performance in the laboratory and in the field has contributed to the slow improvement of crops with improved resistance to abiotic stress in spite of the massive advances in genomic technologies (Bruce et al., 2015). For this reason, our experimental design did not only query the transcriptional response of rice to heat and water deficit in a controlled environment, but also in an agricultural setting. Because field grown plants must incorporate numerous environmental inputs on multiple time scales, analyses of these plants revealed a greater proportion of the underlying regulatory network. However, identifying the underlying environmental change contributing to these regulatory responses is challenging. By including both types of experiments in our analysis, we were able to leverage the complexity of the agricultural field conditions and the specificity of the controlled experiments to interpret the inferred regulatory network. This was particularly the case for TFs whose activities responded to the water deficit treatment in the controlled experiment, as we had somewhat parallel conditions in the agricultural fields. The rain-fed field became drier over the course of the sampling period, particularly during the dry season, contributing to a water deficit field condition.

With our approach we are able to infer target genes only for TFs that have known binding motifs or otherwise proposed targets and so our present network excludes many genes. While this will have led to gaps and mis-assignments in our network, we can use new data as it becomes available to update our network prior. For example, ATAC-seq may be combined with data of ChIP-seq, DNA methylation patterns, and novel TF binding motifs in more sophisticated ways than shown here (Hoffman et al., 2012). This would increase the resolution of the inferred EGRIN by breaking up predictor groups, as well as reducing the number of indirect interactions that we predict.

The network and source data provided with this work provide a vital starting point for studying and manipulating stress responses in rice. Because gene regulatory networks define plant response to environmental signals, divergence of EGRINs is hypothesized to be an important source of ecotypic diversity (Thompson et al., 2015; Weirauch and Hughes, 2010). In future analyses, it will be possible to incorporate additional genome scale measurements, including variations in the non-coding parts of the genome, and chromatin immunoprecipitation, and by reprogramming regulatory interactions using gene lesions and manipulations at the transcriptional level (Chavez et al., 2015; Konermann et al., 2014; Maeder et al., 2013), to expand the EGRIN. Learning accurate genome scale EGRINs can be used to identify high-priority targets for plant breeding and biotechnology programs, and will permit the study of the evolution of gene networks and their relation to adaptive diversification of plant species in different environments.

## Methods

### Plant materials and growth conditions

Seeds of the five rice landraces used in this experiment were obtained from the International Rice Research Institute (IRRI):Azucena (AZ; IRGC#328, *Japonica*), Kinandang Puti (KP; IRGC#44513, *Indica*), Palawan (PA; IRGC#4020, *Japonica*), Pandan Wangi (PW; IRGC#5834, *Japonica*), and *Tadukan* (TD; IRGC#9804, *Indica*). AZ, KP, PW and TD were used in the controlled growth chamber experiments; AZ, PW, and PA were used in the field experiments. All experiments were conducted at the International Rice Research Institute in Los Banos, Philippines.

Single factor perturbation experiments were conducted in walk-in growth rooms. Seed dormancy was broken by incubating seeds for five days at 50°C in a dry convection oven. Seeds were germinated in water in the dark for 48h at 30°C and were then sown on hydroponic rafts suspended in 1X Peters solution (J.R. Peters Inc., Allentown, PA). The pH of the growth media was maintained at 5-5.5 throughout. Plants were grown for 14 days in climate-controlled growth chambers (12h days; 30°C/20°C day/night, 300-500 μmol quanta m^-2^ s^-1^ at the leaf surface). Relative humidity was maintained between 50-70%. Seeds and plants for the field experiments were treated as described previously (Plessis *et al.* 2015).

### Experimental treatments

Single factor perturbation experiments were conducted on 14-day-old seedlings. Treatments began precisely 2h after the chamber lights were turned on. Samples were collected every 15 minutes for up to 4.5h for each of five treatments:control, heat shock, recovery from of heat shock, water deficit, recovery from water deficit. Heat shock was initiated by moving the hydroponic rafts to a 40°C climate controlled growth chamber (RH and light intensity as above). After 2h, some plants were returned to the 30°C chamber (heat shock recovery); the remainder were kept in the 40°C chamber for the duration (heat shock). Water deficit was initiated by removing the rafts from the hydroponic media and allowing the roots to air-dry. After 1.5h, some of the plants were returned to hydroponic tanks containing Peters solution (recovery from water deficit); the remainder were kept in tanks without water (water deficit). Each sample comprised the leaves - from 10 cm above the seed - of 16 juvenile plants. Samples were harvested and flash frozen in liquid nitrogen for each treatment, time point, landrace, and replicate. The entire experiment was replicated on two consecutive days yielding two biological replicates for each condition. Samples from field experiments were harvested as described previously (Plessis *et al.* 2015). Aliquots of these samples were used for the RNA-seq analyses.

### Gas exchange

Instantaneous photosynthetic rate and stomatal conductance of the youngest fully expanded leaf was measured using a portable gas analyzer (Li-6400; Licor, Lincoln, NE,USA). Conditions in the leaf cuvette were set at a saturating light intensity of 1000 μmol quanta m^-1^s^-1^ with a target air temperature of 30°C (40°C in the case of the heat shock measurements), reference CO_2_ concentration of 400p,mol mol^-1^ and humidity of 70%. The measurements were corrected for leaf area. Time 0 for the heat treatment coincides with minimum functional status observed within 15 min of the start of the stress imposition. Further, the break within the trace coincides with the transition from 40 to 30°C where there was instrument instability.

### RNA extraction and library preparation

All protocols were conducted as per the manufacturers’ instructions. Total RNA was extracted using RNeasy Mini Kits (Qiagen). RNA quality was determined by gel electrophoresis. Contaminating DNA was removed from the total RNA samples by treatment with Baseline-Zero DNase (Epicentre, Madison, WI, USA), and ribosomal RNA was removed using the Ribo-Zero rRNA Depletion Kit (Epicentre, Madison, WI, USA). Strand-specific RNA-seq libraries were synthesized using the Plant Leaf ScriptSeq Complete Kit (Epicentre, Madison, WI, USA). The libraries were sequenced – either six or eight libraries per lane – using standard methods for paired-end 51 base pair reads, on an Illumina HiSeq 2000 at the New York University GenCore facility. RNA-seq summary statistics are provided in Supplemental File S11.

### RNA-seq processing

The reads were aligned to the *O. sativa* Nipponbare release 7 of the MSU Rice Genome Annotation Project reference (Kawahara et al., 2013) which consists of 373,245,519 base pairs of non-overlapping rice genome sequence from the 12 rice chromosomes. Also included are the sequences for chloroplast (134,525 bp), mitochondrion (490,520 bp), Syngenta pseudomolecule (592,136 bp), and the unanchored BAC pseudomolecule (633,585 bp). The annotation contains 56,143 genes (loci), of which 6,457 had additional alternative splicing isoforms resulting in a total of 66,495 transcripts.

We used Tophat (Kim et al., 2013; Trapnell et al., 2009) version 2.0.6 to align the reads, discarding low-quality alignments (quality score below 1). To count the number of reads that uniquely mapped to genes we used HTSeq (Anders et al., 2014) version 0.6.1. We compensated for variable sequencing depth between samples using the median-of-ratios method of DESeq2 (Love et al., 2014) version 1.6.3, and further performed a variance stabilizing transformation provided by the same package. We used the normalized count data for downstream analysis. Replicates were averaged, except for differential expression analysis (see below). We removed all genes that had zero counts in more than 90% of the conditions. We also removed all genes that has a coefficient of variation smaller or equal to 0.05. This left us with 25,499 genes.

### Differential Expression

For the chamber experiments, we determined differentially expressed (DE) genes using DESeq2 and the raw read counts as reported by HTSeq. Replicates were not averaged and for every cultivar we created a group factor in the design matrix that was treatment and time specific, and accessed the contrasts individually (e.g. “Azucena heat 60min” vs “Azucena control 60min“). This design allowed DESeq2 to estimate gene expression dispersion parameters based on all available samples for a given cultivar. For every gene and cultivar-condition combination this gave us a log-fold-change value and an adjusted p-value. To visualize the similarities and differences between the conditions and the cultivars with respect to differentially expressed genes, we applied Kruskal’s non-metric multidimensional scaling to the fold change matrix. Only genes with at least one cultivar-condition where the adjusted p-value was below 0.0001 and the absolute fold-change was greater than 2 were considered (2097 genes). The distance metric was Euclidean.

### ATAC-seq library preparation

We prepared ATAC-seq libraries from leaves at control (30min, 2h, 4h), heat (30min, 2h, 4h), heat-recovery (4h), water deficit (2h), and water deficit-recovery (4h) conditions in an effort to identify the maximum number of open chromatin regions relevant to our experimental conditions. Two biological replicates were prepared for each condition. The second leaves of 2-week old Azucena rice seedlings were harvested and flash frozen in liquid nitrogen. Intact nuclei were isolated using a protocol generously shared with us by Drs. Wenli Zhang and Jiming Jiang (personal communication). Briefly, ground tissue was suspended in nuclear isolation buffer and washed repeatedly using nuclear wash buffer following a standard nuclear isolation protocol. Chromatin was fragmented and tagged following the standard ATAC-seq protocol (Buenrostro et al.,2015). Libraries were purified using Qiagen MinElute columns before sequencing. Libraries were prepared from control, water deficit and heat-treated plants after 0.5, 2, and 4h. Libraries were sequenced as paired-end 51 base pair reads on an Illumina HiSeq 2500 instrument in the RapidRun mode at the New York University GenCore facility.

### ATAC-seq data processing

The average number of reads was 14.8M and the total for all libraries was 266,839,208 reads. We used Bowtie version 2.2.3 to align the reads to the reference genome. The alignment rate was around 92%. For downstream analysis we removed PCR duplicates using samtools rmdup and required alignment quality scores > 30. This step resulted in a marked reduction of reads as many reads originated from redundant regions of the chloroplast genome or from nuclear encoded chloroplast genes. The final number of aligned reads for downstream analysis is 29,312,972 (11% of all reads; 0.078 reads per bp in the reference genome).

To compare the 18 ATAC-seq samples to each other with respect to location and number of ATAC-seq cut sites (first base of an aligned fragment and first base after the fragment), we counted the number of cuts in all non-overlapping windows of 1000 bp in each library. For each pair of libraries, we then calculated Pearson correlations of number of cuts (in log space after adding a pseudo count). Results are shown as Supplementary Figures S5 and S6 illustrating the high reproducibility of the ATAC-seq data, but also how similar the ATAC-seq results are for a given tissue even for samples exposed to different stresses.

In order to define an atlas of accessible regions to be used in network inference we combined the ATAC-seq results from all libraries to maximize the number of identified nucleosome-free regions in the genome relevant to our experimental conditions. To define “open” regions we counted the number of ATAC cut sites that fell into the 72 bp window centered on each base. We considered a base open if its window contained at least one cut site in more than half of the libraries. If two open bases were less than 72 bp apart, we called all intermediate bases open.

### Combining ATAC data and TF motifs as network prior

We used published TF binding motifs and knowledge of open chromatin based on our ATACseq data to generate a prior gene regulatory network for rice. For this, we obtained rice TF binding motifs from the CIS-BP database (Weirauch et al., 2014) dated 2015/05/18, the TRANSFAC Professional database (Matys et al., 2003) version 140805, and manual curation of literature. Since the 5’ UTR upstream gene regions showed a high enrichment for open chromatin, we assumed that *cis*-regulatory elements have the largest effect. To find relevant motif occurrences, we scanned only the open regions of the rice genome (as determined by ATAC-seq) that were also in the promoter region of a gene (-453 to +137 bp with respect to transcription start site). We used FIMO (Grant et al., 2011) to find motif matches with a p-value below 1e-4, keeping only the best (lowest p-value) match per motif-gene pair. For every TF we filtered the associated motif matches by adjusting the p-values using Holm’s method (controlling the Family-wise error rate) and only keeping matches with an adjusted p-value of less than 0.01. Any gene with a motif match in the open part of its promoter was then recorded as a target of the current TF.

We observed that different TFs can be associated with different motifs but yet can have almost identical sets of targets in the network prior. To prevent amplification of these miniscule differences (which stem from redundant motifs) in the downstream analysis, we unified the targets of TFs that were less than 5% different (binary distance) and assigned the union of the targets to both.

### Removing uninformative edges from the network prior (coherence filtering)

To remove uninformative edges from the network prior, we required an overrepresentation of distinct expression patterns among the prior targets of each TF with respect to all expression patterns observed in the data. We grouped all genes into 160 clusters (square root of 25,499, the number of genes in the data set) to define the set of expression patterns in our data. We used hierarchical clustering with average linkage and 1 minus Pearson correlation as distance function. Then, for each TF and each expression cluster, we tested whether the TF targets in the network prior were enriched for the members of the cluster. If the Fisher’s exact test p-value was below 0.05 we marked all TF targets in the network prior that were also members of the cluster as high-confidence targets. The above procedure was repeated for 64 bootstraps of the expression data (conditions were sampled with replacement), and only interactions with a high-confidence frequency (coherence score) of at least 50% were kept for the coherence-filtered network prior.

### Estimating transcription factor activity

Let *X* be the matrix of gene expression values, where rows are genes and columns represent experiments/samples. Let *P* be a matrix of regulatory relationships between TFs (columns) and target genes (rows). The entries in the prior matrix (*P*, the network prior) are members of the set {0, 1}. We set *P_tj_* to one if and only if there is a regulatory interaction between TF *j* and gene *i* in the network prior. Auto-regulatory interactions are always set to zero in *P*. Estimation of TF activities is then based on the following model (Liao *et al,* 2003; Fu *et al,* 2011):

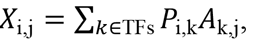

where the expression of gene *i* in sample *j* can be written as the weighted sum of connected TF activities *A*. In matrix notation this can be written as *X* = *PA*, which we solve for the unknown TF activities *A*. This is an overdetermined system, but we can find 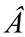 which minimizes 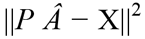 using the pseudoinverse of P. Special treatment is given to time-series experiments, with the modified model:

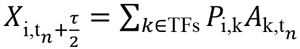

where the expression of gene *i* at time *t_n_* + τ/2 is used to inform the TF activities at time *t_n_*. Here τ is the time shift between TF expression and target expression used when inferring regulatory relationships (see next section). Here we use a smaller time shift of τ*/2*, because changes in TF activities should be temporarily closer to target gene expression changes. If there is no expression measurement at time *t_n_* + τ*/2*, we use linear interpolation to fit the values. In cases where there are no known targets for a TF, we cannot estimate its activity profile, and remove the TF from the set of potential regulators.

### Network inference

The main input to the network inference procedure is the expression data X, the estimated TF activity 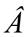, and the known regulatory relationships encoded in the matrix P. The core model is based on the assumption that the expression of a gene *i* at condition *j* can be written as linear combination of the activities of the TFs regulating it. Specifically, in the case of steady-state measurements, we assume

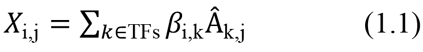

For time-series data, we explicitly model a time-shift between the target gene expression response and the TF activities:

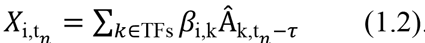

Here, we are modeling the expression of gene *i* at time t_n_ as the sum of activities at time t_n_ – τ, where t_n_ is the time of the n^th^ measurement in the time-series and τ= 15 minutes is the desired time-shift. In cases where we do not have measurements for 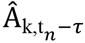 we use linear interpolation to add missing data points.

The goal of our inference procedure is to find a sparse solution to β, i.e. a solution where most entries are zero. The left hand sides of Equation (1.1) and (1.2) are concatenated as response, while the right hand sides are concatenated as design variables. We use our previously described method Bayesian Best Subset Regression (BBSR) (Greenfield *et al*, 2013) to solve for β. With BBSR we compute all possible regression models for a given gene corresponding to the inclusion and exclusion of each potential predictor. For a given target gene *i*, potential predictors are those TFs that have a known regulatory effect on *i*, and the ten TFs with highest mutual information as measured by time-lagged context likelihood of relatedness (CLR) (Greenfield *et al*, 2010; Madar *et al*, 2010). Prior knowledge is incorporated by using a modification of Zellner’s g-prior (Zellner, 1983) to include subjective information on the regression parameters. A g-prior equal to 1.1 was used for the combined network described in this study. Sparsity of our solution is enforced by a model selection step based on the Bayesian Information Criterion (BIC) (Schwarz, 1978).

After model selection is carried out, the output is a matrix of dynamical parameters *β* where each entry corresponds to the direction (i.e. correlated or anticorrelated) and strength (i.e. magnitude) of a regulatory interaction. To further improve inference and become more robust against over-fitting and sampling errors, we employ a bootstrapping strategy. We resample the input conditions with replacement, as well as the prior network, and run model selection on the new input. This procedure is repeated 201 times and the resulting lists of interactions are filtered to only keep those observed in more than 50% of the bootstraps.

### Go enrichment analysis

The reference protein of Oryza Sativa was obtained from the Uniprot Database (www.uniprot.org/proteomes/UP000000763). The April, 2013 release of the Gene Ontology (archive.geneontology.org/full/2013-04-01/) was then queried by these ids, returning all annotations attributed to genes in the Oryza Sativa reference proteome. These annotations were propagated via the true path rule, whereby any protein with an annotation to a GO term also gains annotations for all terms that are parents of the given term, as specified by the GO hierarchy. Lastly, a separate proteome for Oryza Sativa was obtained from the Rice Genome Annotation Project (RGAP), available at:ftp.plantbiology.msu.edu/pub/data/EukaryoticProjects/osativa/annotationdbs/pseudomolecules/version7.0/all.dir/. A blastp was performed, using default parameters, and for each Locus ID from the RGAP, the best-matching Uniprot ID was chosen, and the annotations transferred from that Uniprot ID to the Locus ID. Enrichment analysis of predictor targets was performed using the GOstats R package where all genes present in the network were used as background universe.

Data are available from the Gene Expression Omnibus (www.ncbi.nlm.nih.gov/geo/) under accession numbers GSE73609 (RNA-seq field data), GSE74793 (RNA-seq controlled chamber data), and GSE75794 (ATAC-seq data).Code and supplementary materials are available from the Open Science Foundation osf.io/w6d2n/

## AUTHOR CONTRIBUTIONS

Conceptualization:OW, CH, RB, MP; Methodology:OW, CH; Software:CH, RB; Formal Analysis:CH, RB; Investigations:OW, MMHP, ABN, AP, GMP; Resources:EMS, SVKJ, GBG; Data curation:CH, OW; Writing – Original Draft:OW, CH; Writing – Reviewing & Editing:RB, MP, and all authors; Visualization:CH, OW; Supervision:RB, MP; Project Administration:OW, CH, MP, RB; Funding Acquisition:EMS, RB, MP. The authors declare that they have no competing interests.

## ACKNOWLEDGEMENTS

We thank Ms. Maria Pokrovskii and Dr. Emily Miraldi for their guidance on the ATAC-seq experimental protocol and analysis, Dr. Noah Youngs for sharing his working Gene Ontology annotations with us, Drs. Wenli Zhang and Jiming Jiang for sharing their nuclear isolation protocol with us, Zennia Jean Gonzaga, John Carlos Ignacio, Josefina Mendoza, Eloisa Suiton, and Junrey Amas for their help in preparation of planting, collecting and preparation of leaf samples, and Drs. Zoe Joly-Lopez and Simon Cornelis Groen for thoughtful comments on the manuscript.

## SUPPLEMENTAL INFORMATION

Supplemental Figure S1:Plant functional measurements

Supplemental Figure S2:Distribution of expression values for all genes and control conditions

Supplemental Figure S3:Multi-dimensional scaling plots of differentially expressed genes

Supplemental Figure S4:ATAC-seq fragment size of aligned reads

Supplemental Figure S5:ATAC-seq reproducibility scatter plot

Supplemental Figure S6:ATAC-seq reproducibility heatmap

Supplemental Figure S7:Comparison of ATAC-seq and DNase-seq data in rice

Supplemental Figure S8:TFA stability for all TFs in the network prior

Supplemental Figure S9:Convergence on a stable inferred network

Supplemental Figure S10:Expression of the targets of EPR1 in a circadian data set

Supplemental File S1:Differentially expressed genes

Supplemental File S2:Open chromatin regions

Supplemental File S3:Sample order for heatmaps and TFA plots

Supplemental File S4:The network prior

Supplemental File S5:TF predictor groups

Supplemental File S6:Mean of estimated TF activities

Supplemental File S7:TFA plots

Supplemental File S8:Inferred network

Supplemental File S9:Network predictor statistics

Supplemental File S10:Gene Ontology enrichment results

Supplemental File S11:RNA-seq summary statistics

